# Predicting Anti-Cancer Drug Combination Responses with a Temporal Cell State Network Model

**DOI:** 10.1101/2022.10.05.511031

**Authors:** Deepraj Sarmah, Wesley O Meredith, Ian Weber, Madison R Price, Marc R Birtwistle

## Abstract

Cancer chemotherapy combines multiple drugs, but predicting the effects of drug combinations on cancer cell proliferation remains challenging. We hypothesized that by combining knowledge of single drug dose responses and cell state transition network dynamics, we could predict how a population of cancer cells will respond to drug combinations. We tested this hypothesis here using three targeted inhibitors of different cell cycle states in two different cell lines. We formulated a Markov model to capture temporal cell state transitions between different cell cycle phases, with single drug data constraining how drug doses affect transition rates. This model was able to predict the landscape of all three different pairwise drug combinations across all dose ranges for both cell lines with no additional data. This work shows how currently available or attainable information could be combined to predict how cancer cell populations respond to drug combinations.

## Introduction

Matching chemotherapy regimens to cancer patients remains a grand challenge of oncology and personalized medicine. Targeted drugs often have genetic biomarkers, such as BRAFV600E for vemurafenib^1,2^, EGFR mutations and copy number amplification for gefitinib^3,5,7^, BCR-ABL fusion for imatinib^9,11^, and HER2 copy number amplification for trastuzumab^13,14^. However, such matched patients often do not respond to therapy and/or eventually develop resistance. Why? One major driver is tumor heterogeneity; cells in different “states” that have different drug sensitivities. Cell states are often either defined by their histology or transcriptomics (through for example single cell RNAseq experiments)^15–21^, and it is becoming appreciated that cells can transition between such states in development-like networks, sometimes called cell state networks^17,22^. Such plasticity between cell states can contribute to drug resistance^23–25^, and combinations of drugs targeting different pathways and factors involving phenotype transition have been proposed to prevent such resistance^23^. Another is the multi-variate complexity of biochemical networks within which drug targets reside and by which chemotherapy drugs exert their action^26–31^. These networks can differ between cell states, adapt to therapy, and also give rise to non-intuitive therapy results, such as feedback loops and compensatory pathways underlying the efficacy of combining Raf and MEK inhibitor combinations, which lie in the same genetic pathway^32–35^.

Massive agnostic efforts have screened thousands of cancer cell lines for sensitivity to hundreds of anti-cancer drugs, with matched multi-omic data to mine for biomarkers predictive of drug response^36–43^. These efforts, while substantial, still have not solved the problem of how to match patients to drugs. Moreover, many chemotherapy regimens comprise combinations of 3-4 drugs. Comprehensive experimental exploration of just 2-way drug combinations for hundreds of anti-cancer drugs across a representative cohort is infeasible clinically, and currently unreachable even in cell culture systems.

The inability to obtain an experimental solution to the problem of matching drug combinations to patients has motivated computational modeling approaches. In principle, more comprehensive exploration of drug combination space could be achieved *in silico.* Various computational methods including mechanistic models and machine learning approaches have shown promise in predicting drug combination responses, especially taking into consideration context specific pathology and omics data as well as identifying specific biomarkers and drug-targets ^44–49^. Regardless of the modeling methods being used, there is a widespread focus on using information about biochemical networks to facilitate drug combination response prediction ^31,50–52^. Despite advanced methods being applied to integrate such information into models, building predictive drug combination response models remains an unsolved challenge. Any solution to this problem must invariably rely on experimental data that is already existing or is realistically attainable, such as single drug dose responses.

In this paper, rather than focus on modeling biochemical networks, we test the hypothesis that by combining knowledge of single drug dose responses and cell state transition network dynamics, we could predict how a population of cancer cells will respond to drug combinations. Although this hypothesis runs contrary to the predominant biochemical network-centered view of this problem, cell state transitions are largely governed by biochemical networks in which drug targets are embedded, so in a sense this idea is encompassing prior logic. We test this hypothesis by focusing on three drugs that target cell cycle transitions in two different cell lines. A Markov model is developed to capture population growth and single drug dose responses, and then this model is used to predict all two-way drug combination responses with no further adjustment. Comparison of these model predictions to experimental tests shows surprisingly good agreement. These results suggest a sufficient formulation for predicting how cell population growth dynamics respond to drug combinations that relies on currently available and attainable information, and as such could have widespread impact for precision oncology.

## Results

To test our hypothesis that knowledge of single drug dose responses combined with cell state transition network dynamics could enable prediction of drug combination responses **(Fig. 1a),** a model system is needed. There are a variety of choices for cell state transition networks and drugs which modulate them; here we focus on the cell cycle and three targeted kinase inhibitors (**Fig. 1b**). Specifically, we focus on a MEK1/2 inhibitor (PD0325901) that primarily blocks transition of G_0_/G_1_ cells ^53^, a CDK4/6 inhibitor (abemaciclib) that primarily blocks transition of (late)G_1_S cells ^54–57^, and a PLK1 inhibitor (TAK-960) that primarily blocks transition of G_2_/M cells ^58–61^. Drug dose response experiments evaluating cell number after 3 days of treatment show that both U87 and U251 cells are responsive to these drugs as single agents (**Fig. 1c**).

**Figure 1.**
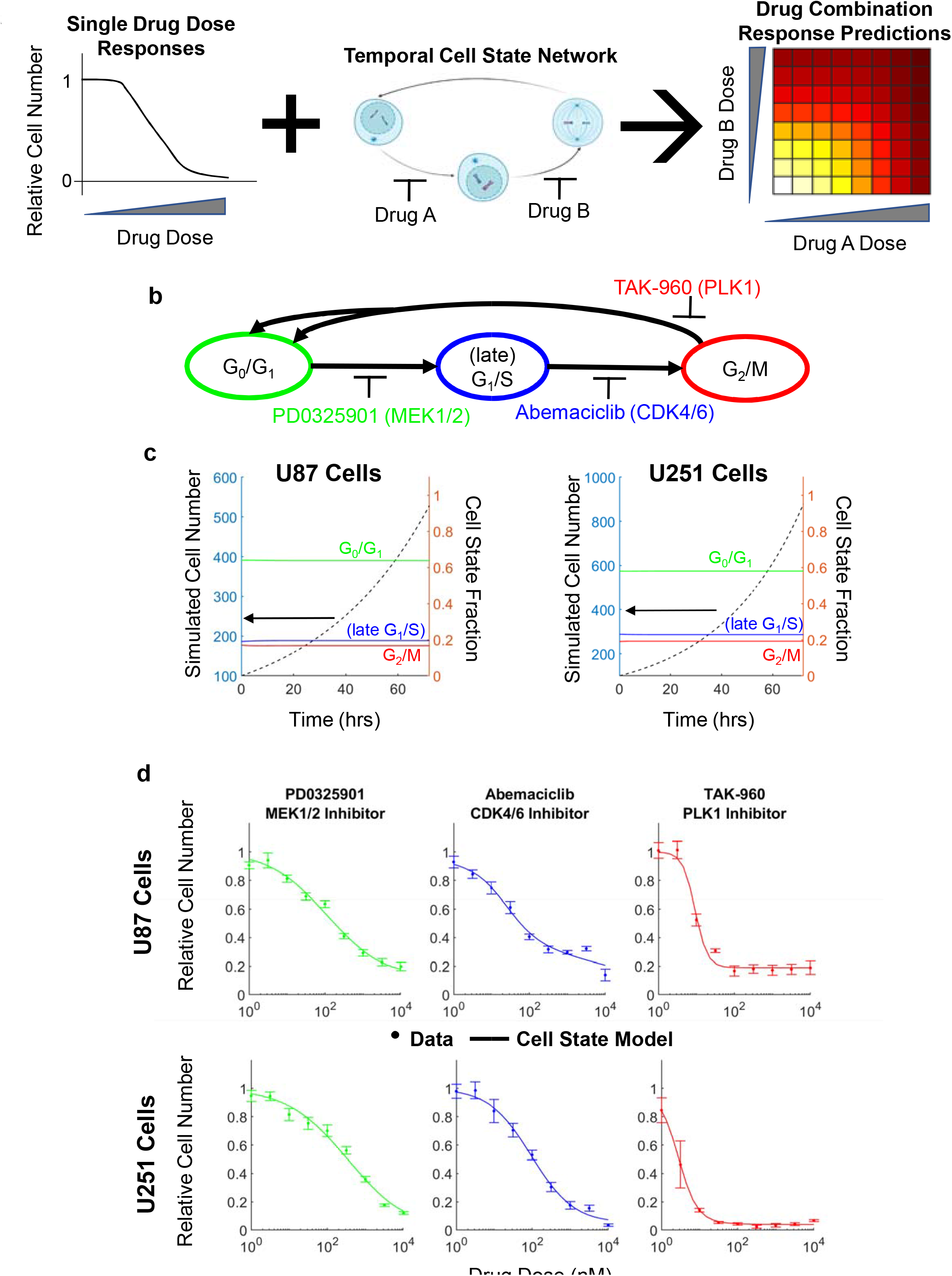
Modeling temporal cell states and single drug dose responses. **(a)** Graphical abstract. Schematic showing that integration of single drug dose response experiments with temporal cell state network can predict drug combination responses **(b)** Schematic of the temporal cell state network comprising G0/G1, late G1/S and G2/M states and the activity of the drugs PD0325901, Abemaciclib and TAK960 in each state **(c)** Time courses of cell number in the temporal cell state model for U87 and U251 cells starting with 100 cells for 72 hours. Cell proportions at G0/G1, late G1/S and G2/M states are also mapped which remain constant. **(d)** Single drug dose responses for PD0325901, Abemaciclib and TAK960 in U87 and U251 cells at 72 hours. The temporal cell state model can capture the single drug dose responses reasonably well.

Before accounting for drug effects, we first constructed and parameterized a temporal cell state network model based on Markov formalisms (see Methods) that describes cell population dynamics in the absence of drug for U87 and U251 cells. Cells in the G_0_/G_1_ state can transition to the (late)G_1_S state, which can then transition to the G_2_/M state. Upon transition from G_2_/M to G0/G1, cell division occurs, increasing cell number by one. For this case without drug, we consider cell death transitions (which decrease cell number by one) to be negligible (we include them below to capture high dose features of some single drug responses). In each time step (chosen to be 1 hr), cells can either remain in their current state, or transition. We set the three unknown transition probabilities for each cell line by requiring agreement with population doubling time (30 hours-U87^62–64^ and 24 hours-U251^65–67^ and the steady-state cell state ratios (64:19:17 for U87^68–72^ and 58:23:19 for U251^73–75^) (**Fig. 1c**). Simulations recapitulate these features.

Drug action was modeled by assuming the transition parameters are a sigmoidal function of drug dose (see Methods). Fitting to the single drug dose response data yielded excellent agreement between model and data (**Fig. 1d**). At high doses for some drugs / cell lines, small cell death transition terms were included to account for the fact that observed cell numbers were lower than the initial number of cells (see Methods). Overall, these results demonstrate that the Markov model of cell state transition dynamics can capture cell population growth and single drug responses for the investigated system.

Now that we had a model that could take as input any of the three drugs at any dose and simulate cell population dynamics, we could predict how drug combinations would affect cell number for all pairwise combinations of the three drugs (**Fig. 2a-b, left**). These predictions demonstrate good agreement with independent experimental data for every drug combination for each cell line (**Fig. 2a-b-right**). Notably, no modifications were made to the model—only information about the cell state transition network dynamics and single drug dose responses were needed to perform this prediction.

**Figure 2.**
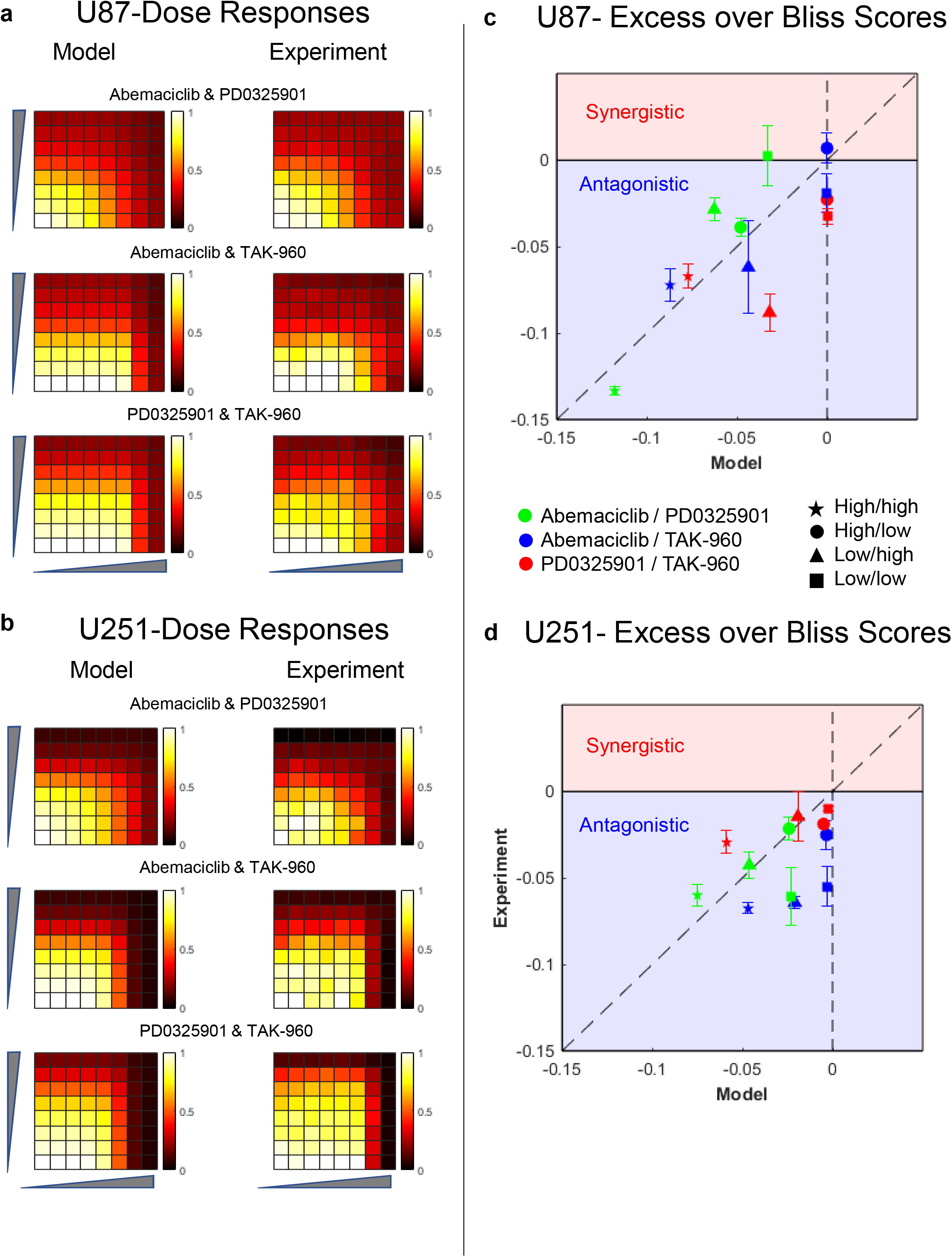
Combination dose responses: Model prediction vs experiments. **(a,b)** Predicted and measured combination drug dose responses for Abemaciclib/PD0325901, Abemaciclib/TAK960 and PD0325901/TAK960 for U87 cells(a) and U251 cells (b) **(c,d)** Experiment vs model excess over Bliss scores for Abemaciclib/PD0325901, Abemaciclib/TAK960 and PD0325901/TAK960 for U87 cells(c) and U251 cells (d). For a pair of drugs(say Drug A/Drug B), the combinations of the lowest 4 doses of both drugs (besides zero doses) are categorized as low/low, combinations of the highest 3 doses of Drug B with the lowest 4 doses of Drug A are categorized as high/low, combinations of the lowest 4 doses of Drug B with the highest 3 doses of Drug A are categorized as low/high, and combinations of the highest 3 doses of both the drugs are categorized as high/high. This is further illustrated in Figure S1.

Analysis of drug combination responses often includes assessment of drug synergy or antagonism, a more qualitative and categorical analysis. A common metric that we use here is excess over Bliss ^76^, which quantifies how much of the observed drug response is beyond statistically independent action by each drug. In particular, we used a variation of excess over Bliss that is more robust and reproducible because it uses sigmoidal fits to data to mitigate the impact of experimental noise in any single data point ^77^. Values close to zero indicate noninteracting combinations, positive values indicate synergistic combinations and negative values indicate antagonistic combinations. We stratified the data and model predictions into four dose quadrants (high/high, low/high, high/low, and low/low) and evaluated agreement between the two (**Fig. 2c-d; Fig. S1**). Both model and experiment show drug combinations were predominantly mildly antagonistic or non-interacting. Overall, these results provide support for the hypothesis that responses to anti-cancer drug combinations can be predicted with a model of cell state transition dynamics and knowledge of single drug dose responses.

## Discussion

Predicting how varied drug combinations control cancer cell population growth is key to improving cancer precision medicine. Experimental solutions alone cannot cover the vast combinatoric space comprising drug combinations and different cancer cell types, necessitating computational approaches. Any computational approach should rely only on data that is available and/or feasibly attainable. Here, we explore the use of a computational approach that, rather than focus on biochemical networks in which drug targets reside, focuses on cell state networks where drugs influence transitions. By combining information about the cell state network dynamics with single cell drug dose responses, we were able to predict combination responses for three different targeted anti-cancer drugs in two different cell lines with no additional model modifications. We expect this finding to be impactful as it informs expansion to different drugs and cell types.

While we do not explicitly consider the role of biochemical networks in drug combination response, in a sense, they are implicitly accounted for in the mapping of drug concentration to cell state transition rates. In the investigated case, there was an arguably clean mapping of drug concentration to single transition rates, which simplified the effort. In other cases, such mapping may not be known *a priori* and/or more complex, i.e., a single drug may influence multiple transition rates. Biochemical network models that capture such complexities or mapping may prove useful in such situations^26–30,45,78^. Assumptions regarding the additivity (or not) of multi-drug action on transition rates would have to be asserted. The current availability of drug combination response data sets ^79–82^could facilitate the testing of such methods. Such future work could explore drug combination features we did not here, such as combining drugs that are not effective as single agents. They could also explore conditions that lead to drug combination synergy; the systems chosen here exhibited predominantly antagonistic behavior. Avoiding antagonism, however, is an important goal. It is thought that a small fraction of all drug combinations lead to synergistic behavior, but finding them, and how synergy is controlled by cell type, is of critical importance for precision oncology.

Application of this approach to other systems requires identification of cell state network models. Again, in our case the cell cycle is well established in terms of structure, but other such networks may not be. Studies have also confirmed the factors behind certain other cell state transitions-for instance, the transcriptomic factors and signaling molecules in different epithelial to mesenchymal transitions^30,83–85^. Cell state transition networks have been identified for multiple cancer types^15–19,21,22,86–88^, generally by combining single cell measurements (e.g. single cell RNAseq), with perturbation time courses, such as enriching for one cell state and then observing the fractional composition dynamics. Recently, we proposed a general theory built upon modular response analysis^89–92^ that allows one to reconstruct cell state networks from such perturbation time course data^78^. This theory is compatible with the Markov formalisms used here. Such Markov formulation may have further applicability to other cell state systems^22,87,88,93–95^, but other approaches have been used^96–99^. Cell state transitions are subject to inherent stochasticity and describing the cell transitions as a Markov process is a common tool to capture this probabilistic aspect. However, this also relies on the assumption that the transition probabilities and the underlying variables are known and that the cell states are properly sampled and well classified. There could be several knowledge gaps in these assumptions including that cells may be transcriptomically intermediate between canonically defined states^100^ and biological data may be sparse^101^. Inclusion of methods such as lineage tracing and methods able to handle sparse data^98^ may help address some of these gaps.

Overall, we have tested a relatively simple hypothesis that knowledge of single drug dose responses combined with cell state network dynamics is sufficient for prediction of drug combination responses. This hypothesis seems to hold true at least for the three drugs and two cell lines studied here, providing a potentially powerful rationale for guiding drug combination response modeling efforts. Expansion to more cell lines, cell state systems, and drugs will of course be important for further testing. Our findings here provide an important step towards being able to predict how cancer cell populations will respond to combinations of anti-cancer drugs, a key capability for cancer precision medicine.

## Data Availability

The code needed to reproduce the data and figures are included and can be accessed at DOI: 10.5281/zenodo.7150564

## Acknowledgments

MRB acknowledges funding from the National Institutes of Health Grants R35GM141891 and support from the Clemson Creative Inquiry program.

## Author Contributions

MRB and DS conceived of the work. DS made the figures. DS, WOM, IW and MRB performed the experiments. DS and MRB wrote the manuscript.

---

**Figure S1:**
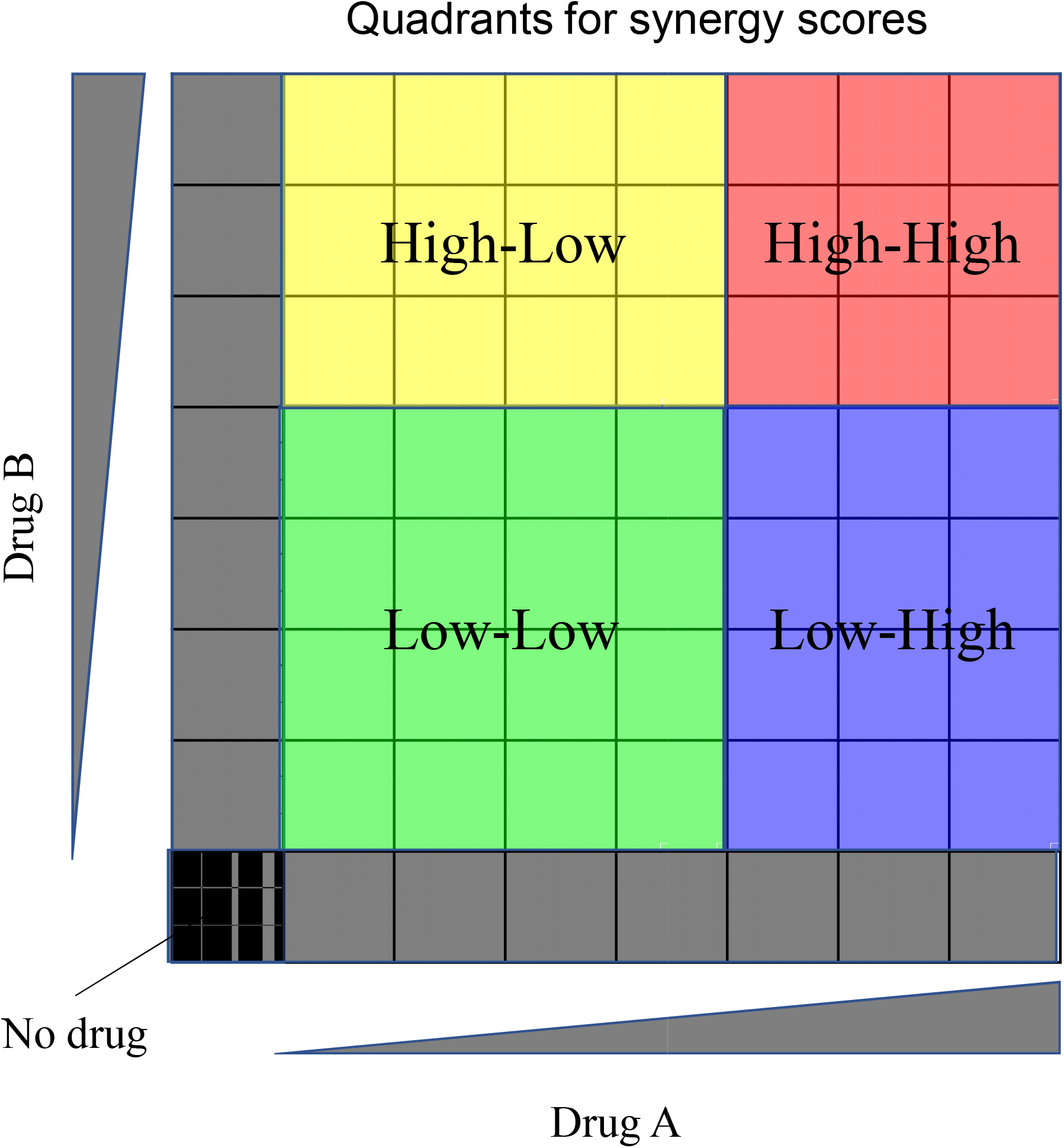
Schematic of combination drug dose matrix for two drugs and quadrants to measure synergy scores across-low/low, high/low, low/high and high/high doses of drugs.

## Methods

### Experimental Methods

#### Cell Culture

U87 and U251 cells (both STR profiled internally) were cultured in full growth medium comprising DMEM (Gibco #10313039) supplemented with 10% FBS (Corning #35-011-CV) and 2 mM L-Glutamine (Corning #25-005-CI). The cells were cultured at 37°C in 5% CO_2_ in a humidified incubator and passaged every 2-3 days with 0.05% trypsin (Corning #25-052-Cl) to maintain sub confluency.

### Drug Dose Response Experiments

U87 and U251 cells were seeded in 96 well plates (Corning-Falcon #353072) with 500 cells per well, counted with a hemocytometer. Cells were seeded in 90 μl full growth media and cultured overnight. The next day, 10 μl of media containing 10X the final drug concentration was added and the plates cultured for 72 hours.

The three drugs were procured from the following sources - PD0325901 (Selleckchem #S1036), Abemaciclib (Selleckchem #S5716) and TAK960 (Tocris #5403). The quantities of each drug-PD0325901 (25 mg, molar mass-482.19g), Abemaciclib (25 mg, molar mass-506.59g) and TAK960 (10 mg, molar mass-598.06g) corresponded to 0.0518 millimoles, 0.0493 millimoles and 0.0167 millimoles respectively and were diluted in 5.18 mls, 4.18 mls and 1.67 mls of sterile filtered DMSO to bring the final concentration to 10 mM for each drug. These dilutions were then aliquoted into 10 μL batches. Before adding to cells, 990μL of full growth media was added to a 10 μL drug aliquot, diluting it to 100 μM, or 10X times the highest desired dose. This concentration was further serially diluted 8 more times in full growth media containing 1% DMSO (to maintain the same DMSO concentration in each dilution) and by a factor of 3.16 each time. This results in 9 dilutions with the drug concentrations between 10 nM-100 μM. In 9 wells with cells seeded overnight in 90 μL media, 10 μL of the serially diluted drugs are added. In the 10^th^ well, 10 ul of full growth media containing 1% DMSO was added as the vehicle control dose.

For combination dose responses, U87 and U251 cells were seeded in 96 well plates (Corning-Falcon #353072) with 500 cells in each well. Eight by eight wells were seeded in 150 μl full growth media and cultured overnight. The next day, 25 μl of media was added twice to each well, each containing 8x of the final drug concentration and cultured for 72 hours. The final drug concentrations were chosen to reflect their responsive range for the cell lines (1.22 nM-5μM for PD0325901 and Abemaciclib, 0.0122nM-50nM for TAK960). Before adding to cells, full growth media was added to a 10 μL drug aliquot, diluting it to 8X times the highest desired dose. This concentration was further serially diluted 6 more times in full growth media containing 1% DMSO (to maintain the same DMSO concentration in each dilution) and by a factor of 4 each time. This results in 7 dilutions with the desired drug concentrations.

### Staining and Computational Image Analysis

After 72 hours of treatment with the drugs, the cells were stained with Hoechst (BDBiosciences #BD 561908) and Propidium Iodide (Millipore Sigma #P4170) at a final concentration of 1 μg/ml and 2 μg/ml to stain all cells and dead cells respectively. After 30 minutes, the wells were imaged using the TagBFP (Excitation-390nm, Emission-447nm) and RFP filters (Excitation-531/40 nm, Emission-593/40 nm) in Cytation 5 (Biotek). Each image was flatfield corrected and background subtracted using CellProfiler. The nuclei were then identified using the IdentifyPrimaryObjects feature, and a pseudo image was generated. The number of all counts of cell nuclei stained with Hoechst and Propidium Iodide were compiled by CellProfiler and exported as csv files. The Propidium Iodide stained nuclei counts were subtracted from the Hoechst stained nuclei counts for each well, and this was taken as the live cell counts.

### Model and Computational Methods

#### A Markov Model of Temporal Cell State Transitions

Consider a Markov state model comprising three nodes, representing cell states G_0_/G_1_, late G_1_/S and G_2_/M, taken as 1, 2, and 3 in short. *M_1_, M_2_* and *M_3_* are the proportion of cells transitioning from states 1-2, 2-3 and 3-1 respectively, within a given timestep-one hour for our considerations. *M_11_, M_22_* and *M_33_* are the proportion of cells that did not transition from states 1, 2 and 3 respectively. A cell in state 3 undergoes cell division which gives rise to two cells in state 1. We formulate this scenario using a jump Markov process model as follows:

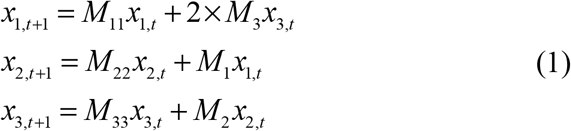

where, *x_i,t_*, are the numbers of cells in state *i* at time point *t.* The sum of cell numbers in each state at a particular time gives the total number of cells at that time. We set the time interval between two Markov jumps at 1 hour and simulate the model for a total of 72 hours.

These equations are subject to the constraints that the proportion of cells within a state must add to 1. Therefore-

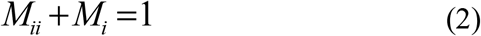

Or

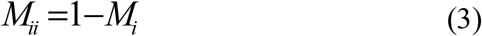

Incorporating this in the above equation enables us to represent the system in terms of *M_1_, M_2_* and *M_3_*

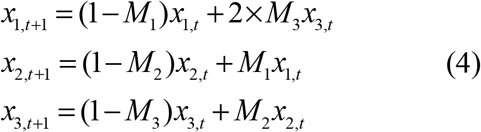

In short this may be represented as follows-

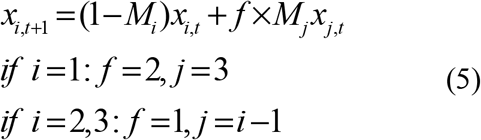

We estimated the unknown transition rate parameters based on experimental data for cell doubling times of 30 and 24 hours and cell state ratios of 64:19:17 and 58:23:19 for U87 and U251 cells respectively (data references in Results). We used fmincon in MATLAB with a least squares formulation giving values of *M_1_, M_2_* and *M_3_* as 0.05 hr^-1^, 0.14 hr^-1^, 0.14 hr^-1^ for U87 cells and 0.075 hr^-1^, 0.16 hr^-1^, 0.16 hr^-1^ for U251 cells. We repeated this estimation 5 times with random initial guesses, each converging to the same values, demonstrating uniqueness of the estimates.

#### Single Drug Dose Response Modeling

Each of the drugs-PD0325901 (MEK1/2 inhibitor), Abemaciclib (CDK4/6 inhibitor), TAK960 (PLK1 inhibitor) were modeled as having an inhibitory effect on the respective Markov parameters *M_1_, M_2_* and *M_3_*. We used a sigmoidal hill-type function to describe the effect of drug concentration on transition rates as follows

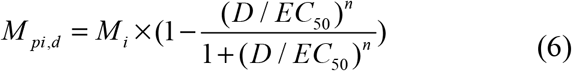

Where, *M_pi,d_* is the perturbed/inhibited Markov parameter *M_i_* after inhibition with the respective drug of dose *D.* The remaining parameters in the equation-half maximal ‘*EC_50_*’ and hill coefficient ‘*n*’ were initially taken as unknowns and were estimated by fitting the respective drug dose response for each drug. We used fmincon in MATLAB to obtain the set of parameters that minimized the sum of squares error relative to the data.

This fit model could explain the drug dose responses reasonably well. However, in some cases, the model fits at higher doses of the drugs were higher than the experimentally obtained cell numbers. The experimentally observed cell numbers at some high doses were lower than the initial number of cells loaded, indicating some cell death. Therefore, an additional parameter *M_φ,i,d_* was introduced for each drug, with a value of zero for no drug and a higher value at greater doses of the respective drug

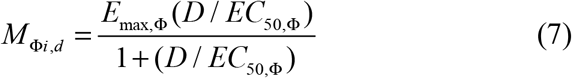

where, *E_max,φ_* is the maximum cell death possible by a high dose of the drug and *EC_50,φ_* is the half maximal drug dose related to cell death.

Eq.5 then changes as follows.

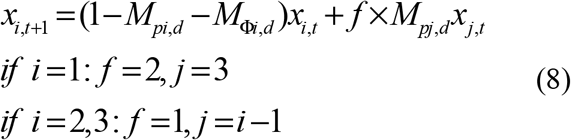

The respective parameters obtained to fit the single dose response parameters for U87 and U251 cells are as follows-

**Table #1.**

| U87 cells | PD0325901 | Abemaciclib | TAK-960 |
| --- | --- | --- | --- |
| $M$ | 0.05 | 0.14 | 0.14 |
| $EC_{50} (\mu M)$ | 0.0769 | 0.011 | 0.0074 |
| $n$ | 0.609 | 0.686 | 2.67 |
| $E_{max}$ | 0.086 | 0.0012 | $3.9 \times 10^{-6}$ |
| $EC_{50,\phi}(\mu M)$ | 1.514 | 0.0274 | 662.4 |

**Table #4.**

| U251 cells | PD0325901 | Abemaciclib | TAK-960 |
| --- | --- | --- | --- |
| $M$ | 0.075 | 0.16 | 0.16 |
| $EC_{50}(\mu M)$ | 0.268 | 0.121 | 0.0024 |
| $n$ | 0.565 | 0.655 | 1.663 |
| $E_{max}$ | 0.016 | 0.022 | 0.018 |
| $EC_{50,\phi}(\mu M)$ | 5.7 | 0.098 | 0.009 |

#### Combination Drug Dose Modeling and Synergy Score

To model combination drug effects for two drugs, drug effects at evaluated doses were simulated together using the above Markov models. In order to quantify combination response in both the model and experimental data, we used a robustexcess over Bliss^77^. Consider each row in the combination drug dose response matrix. One of the drug’s doses would be constant across the row (Drug#1, Fig S1a) but the dose of the other drug (Drug#2, Fig S1a) increases from left to right. This can be captured by a 4-parameter logistic model^77,102^-

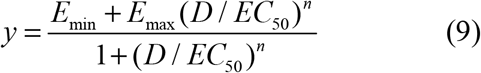

where, *y* is the inhibition effect which in this case is 1 minus the relative cell number with respect to the control*, E_min_* is the minimal possible inhibition effect, *E_max_* is the maximal possible inhibition effect, *EC_50_* is the half maximal drug dose and *n* is the Hill coefficient.

We used `lsqcurvefit` in MATLAB to obtain least-squares estimates for the four-parameters for each of the 8 rows and 8 columns. For a particular drug combination point, the average of the two fitted inhibition values across it’s row and column was taken as the final fitted inhibition value (*y_AB_*).Consider the fitted inhibition values for particular doses of drug A alone (*y_A_*), drug B alone (*y_B_*) and their combination (*y_AB_*). The Bliss independence scores ^37,76,77^ are calculated by

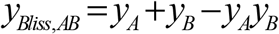

and the excess over Bliss *(EOB)* scores are calculated by

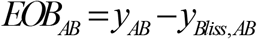

These excess over Bliss scores are calculated for all the three drug combination model predictions for both U87 and U251 cell lines. This exercise is also performed for each of the individual triplicates of the corresponding experimental data.

In order to compare the excess over Bliss scores in the model vs. experiments, the drug combinations are divided into four quadrants (Fig S1a) and the average *EOB* scores across each quadrant are plotted for experimental data vs model predictions (Fig 2c,d).

